# Single Root hair growth under constant force: insights into wall mechanics

**DOI:** 10.1101/2025.10.09.681433

**Authors:** Thomas Alline, Léa Cascaro, Pauline Durand-Smet, Etienne Couturier, David Pereira, Atef Asnacios

## Abstract

Tip growth is a tightly regulated process that enables root hairs to explore their surroundings, enhancing plant development, particularly by improving nutrient uptake. While Lockhart’s viscoplastic framework is widely used to describe this process, it has received limited experimental validation. By integrating optical microscopy with a custom microplate-based rheometer, we created a novel protocol to simultaneously measure, for individual growing root hairs, both the reduction in growth rate and the instantaneous compression in response to a step in applied axial force. The observed growth rate reduction aligns remarkably with a 1D Lockhart viscoplastic model, experimentally validating this framework in tip-growing cells. Additionally, the instantaneous compression upon force application provided an in situ estimate of turgor pressure. Together, these measurements allowed us to determine, for the first time in *Arabidopsis* root hairs, two critical parameters: the yield turgor pressure and cell wall viscosity. Our approach—including the technique, protocol, and analytical framework—can be readily adapted to other tip-growing species and diverse experimental conditions (e.g., varying nutrient availability or osmotic stress). This opens new opportunities to explore cell wall mechanosensitivity and its role in adapting tip growth to environmental signals.

**Significance Statement:** Plant growth relies on their ability to anchor roots in soil and maximize nutrient uptake. This process partly depends on root hairs. These long tubular extensions develop from the root surface, exhibiting a highly directional growth process —tip growth. Understanding how root hair growth adapts to soil mechanics is crucial, especially with climate change and soil hardening.

We present a novel, non-invasive technique to probe the mechanics of root hair walls—key to their growth. By applying a feedback-controlled force, we can investigate the effect of mechanical resistance on root hair growth while preventing buckling, thus accessing elusive cell wall features. This method holds broader significance, as tip-driven growth is also used by fungi and yeasts to colonize their environments.

## Introduction

Colonizing the world is one of the great challenges of living organisms. Animals move themselves, but other organisms, such as plants and fungi, rely mainly on growth. Concentrating the elongation in the tip is a solution for an invasive lifestyle(1), to follow cues (hormonal, chemical, …) while adapting the growth quickly when encountering an obstacle(2). Therefore, tip growth is a convergent evolution observed for cells as distant in the phylogeny as fission yeast(3), fungal hyphae(4), plant pollen tube(5) or plant root hair(6). These cells go through similar phases when they encounter an obstacle: they first slow down and then reorient either actively(7, 8)or through buckling(3); they can pass very narrow gaps without damaging their cellular content(9). The invasive ability is often related to the maximal force these cells can generate, which has been estimated either by deforming calibrated objects, cavities or gaps(1, 3, 8) or with force sensors(7). However, experimental techniques to determine how these forces and the growth process are adapted to environmental cues, including mechanical resistance, are still missing.

Here, we focus on the tip growth of root hairs (RH), which are long cylindrical structures that develop from the surface of differentiated root epidermal cells called trichoblasts. RH explore the soils to improve plant nutrient absorption by increasing the contact surface between the root and its environment(10). They also contribute to the root anchoring in the soil(11). During their development, tip-growing cells have to penetrate a mechanically heterogeneous medium opposing a wide range of forces(12–14).

Plant cell growth originates in an actively regulated osmotic gradient between the cell and the external medium. The entrance of water is counterbalanced by the cell wall tension, generating a hydrostatic pressure called turgor. In turn, cell wall tension positively regulates cell wall growth, leading to a flow law similar to that of a liquid with a stress threshold, namely Lockhart’s law(5, 15). Soil compressive interactions tend to diminish the cell wall tension and reduce the growth rate according to Lockhart’s law(16). Thus, measurement of Lockhart’s growth parameters and their adaptation to external cues is appealing. However, while a considerable amount of theoretical literature was devoted to understanding how Lockhart’s law and other cellular factors (water fluxes, polarity …) are coupled to explain tip growth(17–19), there is still a need for more experimental investigations. Even for the most studied tip growth model, the pollen tube, experimental evidence is limited and sometimes contradicts Lockhart’s law(20). In particular, the pressure threshold for wall flow at the tip, as well as the extensibility (the inverse of the apparent viscosity of the flowing wall), have never been experimentally estimated(5). This is probably due to technical limitations, in particular to the fact that single cells are too small for pressure probes(21) and extensiometry(22), the two traditional methods of Lockhart’s parameter evaluation.

Thus, we designed a new technique allowing us to get strong estimates of Lockhart’s parameters by applying a constant compressive axial force on the tip of a single growing root hair. Compression tests were not traditionally favoured due to the quick buckling of cylindrical structure and the variability of the force pattern (See the External-force method in (23). Nevertheless, techniques coming from classical rheometry, using feedback loops, allow one to apply constant compressive forces on biological samples (24, 25), in particular maintaining them below the buckling threshold for plants(26). Compression tests are attractive as they avoid clamping artefacts associated with extensiometry or pressure probe. In a previous study(27), we explored the elastic behaviour of a RH growing against an obstacle of variable stiffness and we showed the potential of such approaches in estimating Lockhart’s parameters by providing first measurements of *Arabidopsis* RH apparent axial stiffness and excess pressure beyond yield threshold. Here, by applying a controlled compressive axial force on a growing RH, we provide a characterization of growth in a non-invasive manner. Since the applied force is maintained constant during RH growth, we were able to discard the contribution of RH elastic deformations(27) to the measured RH elongation rates. Maintaining the force constant also allows us to average the growth rate temporarily and thus to reduce the noise on the estimated parameters, allowing us to observe a clear linear decrease in the rate of RH growth as a function of the applied force, in line with Lockhart’s prediction. Measuring the rate of RH growth as function of the applied constant force, we could get non-invasive *in vivo* robust estimates of both the excess turgor over the yield threshold and the cell wall effective viscosity of growing single root hairs. Moreover, by analysing the instantaneous compression of the RH in response to a jump in the applied force, we were able to estimate the turgor pressure, and thus the pressure yield threshold. The technique and protocol designed here can be readily used on other tip growing species, and in different conditions (for instance for variable osmotic pressures to mimic the effect of drought), allowing one to get insights in tip growth adaptation to changing environments.

## Results

### A root hair growing under constant force: a Lockhart-like viscoplastic model for root hair tip growth

Here, we adapt to root hair growth a viscoplastic model(3) previously developed to describe yeast growth in confining wells, and we assume that water fluxes are not a limiting factor for growth (see supplementary materials). Turgor pressure generates stresses within the cell wall and, above a critical stress value (the yield stress *Y*_*σ*_), the cell wall is deformed irreversibly with a strain rate proportional to the excess stress above *Y*_*σ*_. Turgor is supposed to be constant during the whole experiment duration (approx. 10 min), and we consider RH growth under constant external force. Since force is kept constant during root hair growth, there is no elastic contribution to changes in root hair length, except for the very brief period immediately following the application of force (<4 s). This observation justifies the use of a viscoplastic model to describe root hair growth, specifically in terms of cell wall flow and elongation. In these conditions, assuming that growth occurs only at the tip in a region of size *l* (figure 1), the rate of root hair growth writes (please see Supplementary Material for details):

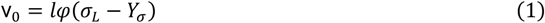

**Figure 1.**
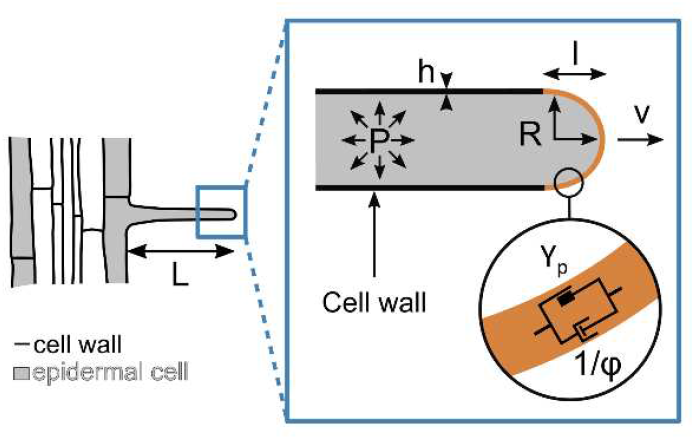
Schematic representing the viscoplastic growth model parameters. The turgor pressure P drive the growth at a speed v. Only a region of size l grows at the tip of the root hair. The root hair radius is noted R and the thickness of the cell wall is denoted h. Y_p_, the yield pressure, and ϕ, the extensibility, are the physical parameters characterizing the wall flow at the RH tip under turgor generated stress in the viscoplastic growth model. These parameters thus characterize wall rheology, Y_p_ representing the threshold in pressure over which the well begins to flow, while 1/ϕ represents the effective viscosity of the flowing wall at the RH tip.

Where *φ* is the cell wall extensibility which can be interpreted as the inverse of a viscosity. *σ*_*L*_ is the longitudinal stress generated in the wall by turgor pressure. Indeed, equation (1) can also be formulated as a function of turgor pressure to get the Lockhart’s law, which states that the growth is proportional to turgor above a yield pressure 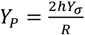:

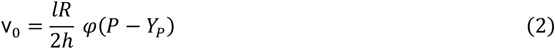

With R the radius of the root hair and h the thickness of its wall (Figure 1).

This model can be easily adapted to describe the effect of an externally applied force at the apex of the root hair. Under the assumption that neither the yield stress nor the extensibility of the root hair depends on the applied force, such a resisting force will decrease the stress in the cell wall, thus reducing growth in a linear manner:

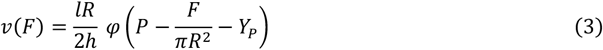

Building on this relationship, we designed a constant force experiment allowing us to determine the extensibility *φ* and the excess pressure (*P* − *Y*_*P*_).

### Quantifying the effect of a constant axial force on root hair growth

To quantify the effect of an opposing force on the rate of root hair growth, we implemented a constant force experiment inspired by a technique we previously designed for single animal cells(25, 28)(Fig 2). Arabidopsis roots were grown on ½ MS agar medium with a microfluidic-like system that have been previously described(29). This allowed us to image the root and root hair cells under a microscope. Before the experiment, we removed some agar gel to make the root hair cell free to grow in liquid MS medium (see the experimental configuration in Fig 2.C). This way, the root was mechanically well maintained to avoid drift upon force application. To apply forces at the growing root hair tip, we designed a specific flexible glass microplate. This microplate acts as a cantilever of calibrated stiffness the deflection of which sets the force applied to the root hair tip. A feedback loop was used to maintain a constant deflection of the microplate, and thus a constant force on the root hair tip. Indeed, during the experiment, a position-sensitive detector (PSD) recorded the cantilever tip position, and this position was maintained throughout the experiment with a feedback loop acting on the piezoelectric stage holding the root. At the beginning of the experiment, a deflection was applied to the microplate, setting the level of the force applied at the RH tip. As the root hair grew, to avoid any further deflection of the cantilever (and thus a change in the applied force), the feedback loop compensates root hair elongation by moving the piezoelectric stage away from the cantilever and with it, the whole root. This maintained the cantilever deflection constant, thus ensuring root hair growth under constant applied force. The root hair growth speed is measured by monitoring the piezoelectric stage displacement necessary to maintain the position of the cantilever tip constant (figure 3.A). The initial root hair growth speed v_0_ was estimated, before force application, for at least 100s. At *t* = *t*_0_ the force applied is set to 1.25, 2.5 or 3.75 µN (figure 3.A-B). Upon force application, we observed an instantaneous compression of the RH of about a µm for the highest applied force, followed by a slower deformation of a fraction of a micron in few tens of seconds (figure 3.B). A linear growth speed resumed after approximately 40s. After force application, the growth speed v(*F*) of the root hair is estimated at *t* = *t*_0_ + 40 *s* during 100s (figure 3.A-B) (orange dashed-line).

**Figure 2.**
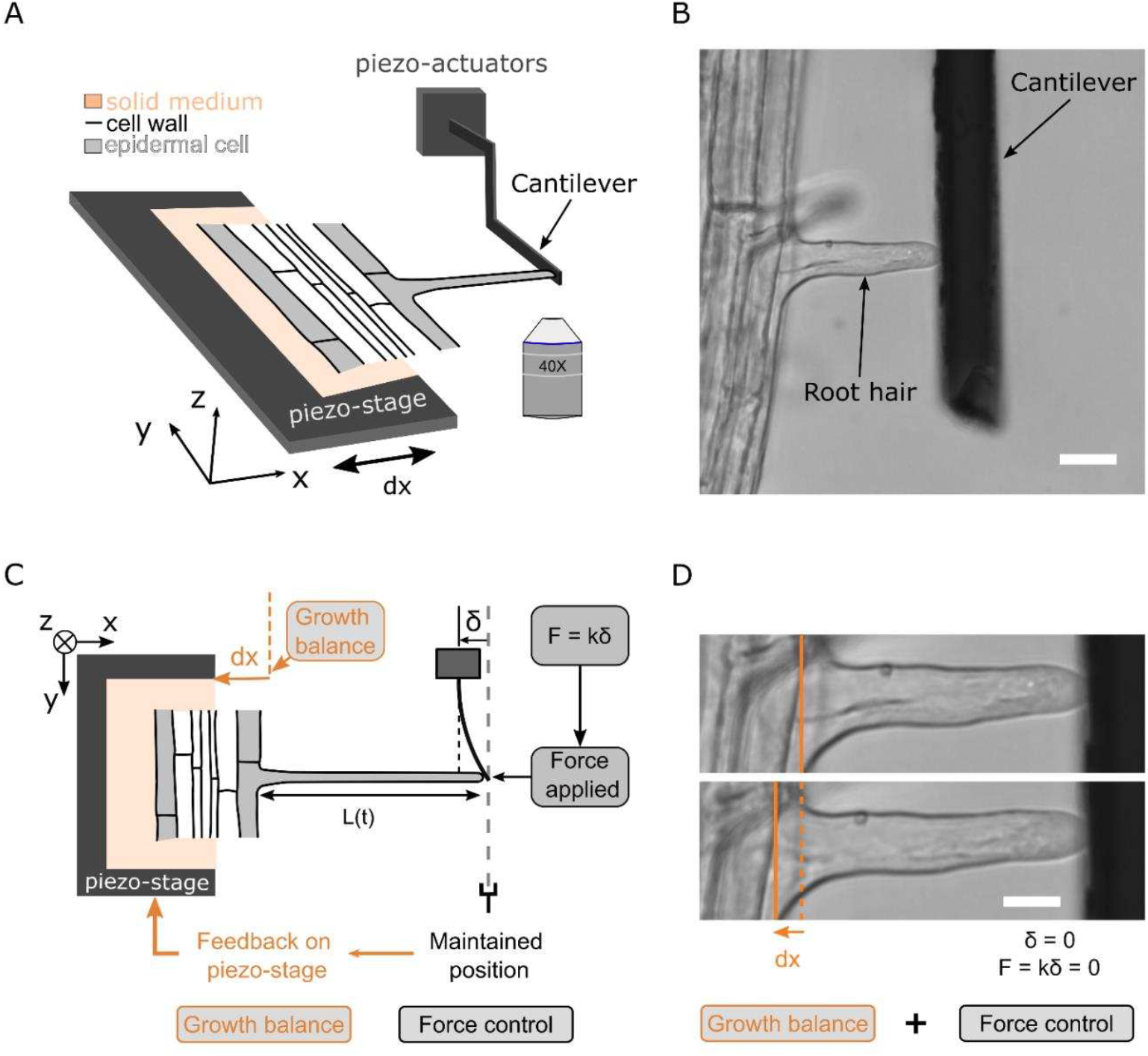
Constant force experiment. **A**. Picture of the experimental system, showing the piezo actuator, the piezo-stage and the glass microplate (cantilever). **B**. Bright field image of the experiment, showing an *Arabidopsis thaliana* root hair growing against the glass microplate (cantilever) (Scale bar = 20µm). **C**. Schematic representation of the feedback allowing to apply a constant force. The optical detector records the position of the contact between the microplate and the tip of the cell. The feedback loop modulates the piezo-stage position dx to keep this position constant when the cell grows against the flexible microplate. **D**. Two bright field images of the same root hair growing against a microplate during a constant force experiment. The orange lines depict the displacement of the piezo-stage. (Scale bar = 10µm).

**Figure 3.**
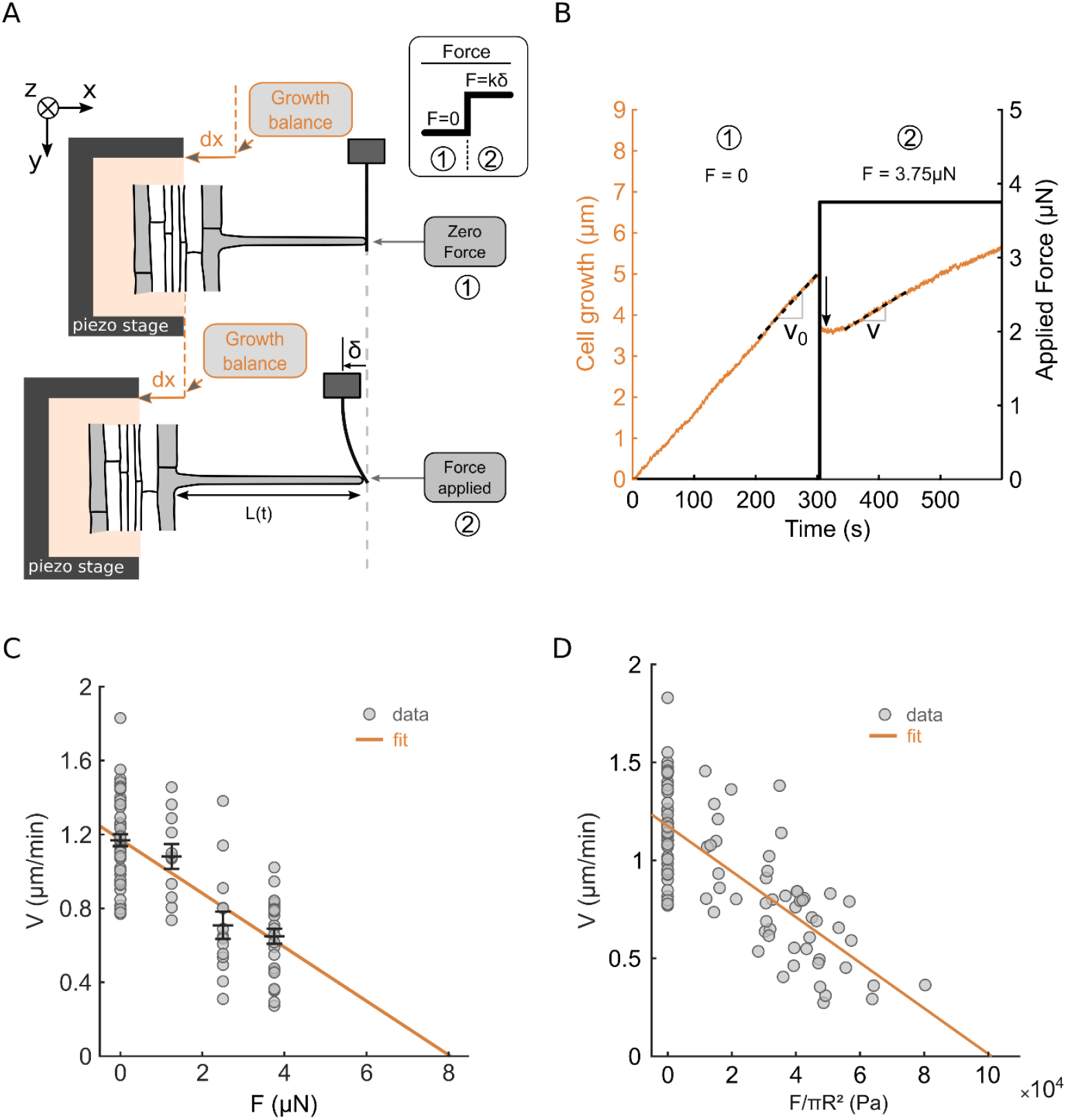
Growth reduction upon force application. **A**. Schematic of the successive step of force application during the constant force experiment. **B**. Representative example of the growth of a root hair during the experiment. The black dotted lines represent the fitted growth rates, 0-100s before force application and from 40-140s after force application. The black arrow refers to the instantaneous elastic RH compression upon force application. **C**. Growth rate as a function of the exerted force. The black error bars represent the average +/-sem for each force exerted. For the 0 nN force point the growth rate before force application is plotted. The orange line corresponds to a linear fit of all data points (n=51 root hairs, from N=13 different plants). **D**. Growth rate as a function of the effective pressure decrease. The orange line corresponds to a linear fit of all data points (n=51, N=13).

The growth speed decreases as the applied force increases (figure 3C), with the free growing rate v(*F* = 0) = v_0_ being 1.17 +/-0.03 µm/min (n=51), while the growth rate was measured at 1.08 +/-0.02 µm/min (n=11), 0.71 +/-0.04 µm/min (n=14) and 0.65 +/-0.03 µm/min (n=26), for respectively 1.25 µN, 2.5 µN and 3.75 µN (figure 3.C). Fitting the data and retrieving the coordinates of the intercept with the force axis, we could also estimate the force needed to arrest RH growth at about 8.0 µN +/-0.65 µN, in the same order of magnitude as the values observes for Yeast(3) and pollen tubes(7). In addition, the length of the root hair does not affect the growth decrease upon force application (figure S1).

To further compare our results to the Lockhart’s model, one needs to represent the growth rate as function of the effective pressure decrease 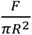 due to the externally applied force (Equation 3). In other words, speed reduction does not only depend on force, but also on RH radius. The radius of each RH tested was measured on bright field images, at 10 µm from the root hair tip at the start of the experiment (distribution in figure S2). The figure 3.D shows the growth rate as a function of 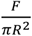. As predicted by the viscoplastic growth model, the growth rate decreased linearly with 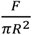 (Equation 3). Adjusting the experimental data with the function given by equation 3 allows one to get a first estimation of the factor 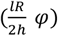 (the slope of the curve), as well as of the excess growth pressure *P* − *Y*_*p*_ (the x-intercept at zero velocity). For instance, we found an excess pressure of *P* − *Y*_*p*_ = 0.101 *MPa* +/-0.018 *MPa*. This value is very close to the first estimation we previously obtained following root hair growth against obstacles of constant or variable stiffness(27). Moreover, this value is close to the one used in tip growth models to ensure consistency with kinetics of pollen tubes’ growth(5).

However, getting the extensibility (*φ*) and the excess pressure (*P* − *Y*_*p*_) from the values of the slope and the x-intercept at zero velocity of a linear fit is not an ideal procedure. First, *φ* and (*P* − *Y*_*p*_) estimations are then related through the fit. Moreover, the estimation of *φ* depends on the values of *l, h* and *R*. And, while *R* is measured for each root hair, using the slope 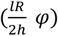 imposes to rely on the mean value of *R* over the population of RH tested, leading to more uncertainties. We therefore defined an alternative data analysis method, enabling us to obtain independent estimates of *φ* and (*P* − *Y*_*p*_), and to reduce uncertainties by accounting, in both cases, for the radius value of each root hair.

### *In vivo* independent estimations of the Pressure above yield and cell wall extensibility of A. thaliana root hairs

Since, for each root hair tested, we measure the rates of growth before (v_0_) and after (v(*F*)) force application, we could compute their ratio v(*F*)/v_0_ as well as the absolute rate reduction v_0_ − v(*F*) allowing us to get independent estimations of the excess pressure (*P* − *Y*_*p*_) and the extensibility *φ* using :

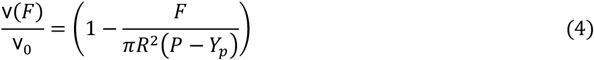

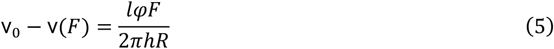

Equations (4) and (5) are retrieved from equation (3).

v(*F*)/v_0_ decreases linearly with the pressure 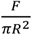 (Figure 4.A). Fitting the data according to equation (4) gives *P* − *Y*_*p*_ = 0.102 *MPa* +/-0.007 *MPa*. This value is very close to the previous estimation from fitting equation (3) to the data of Figure 3.D, and the uncertainty on the estimate is greatly reduced.

**Figure 4.**
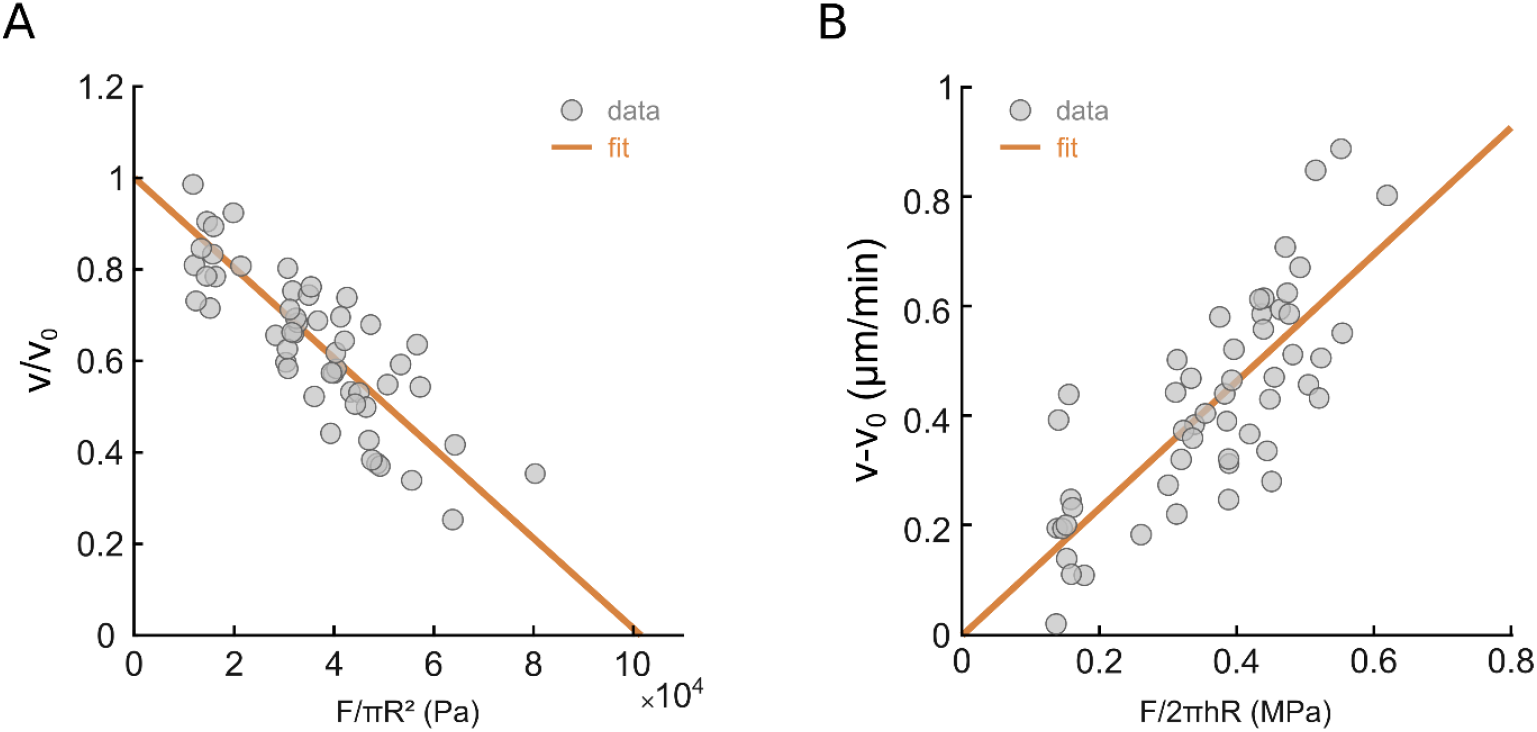
Description of the growth reduction using the viscoplastic model. **A**. Relative growth rate reduction, v(*F*)/v_0_, after force application. The orange line corresponds to a linear fit corresponding to equation (4) (n=51, N=13). **B**. Growth rate reduction, v_0_ − v(*F*), after force application. The orange line corresponds to a linear fit corresponding to equation (5) (n=51, N=13).

Regarding the estimation of *φ*, the viscoplastic model also predicts a linear behaviour of the growth rate reduction v_0_ − v(*F*) as a function of 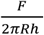, the effective stress reduction in the cell wall upon force application (equation 5). We thus represented v_0_ − v(*F*) as a function of 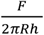 in figure 4.B. Based on the literature(30) we took a cell wall thickness of *h* = 250 *nm*. Fitting Equation (5) to the data of Figure 4.B allows us to estimate the factor *lφ* = 1.93. 10^−14^ *m s*^−1^*Pa*^−1^ +/-0.14.10^−14^ *m s*^−1^*Pa*^−1^. The length of the growth zone has never been measured for the RH of A. thaliana. However, in another experimental model, the root hair of *Medicago trunculata*, a study showed that most of the tip growth occurs in less than 5 µm from the RH tip(6). Assuming this holds true for the root hair of *A. thaliana* (*l* = 5 µ*m*) we can estimate the extensibility of the cell wall at *φ* = 3.8. 10^−9^*s*^−1^*Pa*^−1^ +/-0.28. 10^−9^*s*^−1^*Pa*^−1^.

### *In vivo* estimation of turgor pressure in A. thaliana root hairs trough axial force application

When a force is applied, an instantaneous axial compression is observed (indicated by a black arrow in Figure 3B). To quantify this compression, the total reduction in RH length was measured from brightfield images taken 4 seconds after force application (corresponding to 2 frames in the brightfield image sequence). This measurement allowed us to relate the observed axial compression *dL* to the applied force *F* (Figure 5A). From this relationship, we estimated the axial stiffness of the RH as 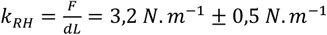. This stiffness value reflects the compression of the entire RH and aligns with previous measurements, which were obtained by monitoring the RH elongation rate against an obstacle of variable stiffness (27). We found that *k*_*RH*_ was one order of magnitude lower than the value expected for elastic compression of the RH wall. Therefore, we attributed the observed stiffness solely to tip compression(27). Indeed, the tip wall can be described as a liquid sheet flowing under the surface stress generated by the difference in hydrostatic pressure. When a force is applied to the tip of the RH, the contact area *S* between the RH and the obstacle increases. By approximating the tip shape as a spherical cap, we used a Hertz-like model to calculate the increase in contact area *dS* from the tip indentation, which we assumed to be equal to *dL*. Under these assumptions, the RH tip behaves like a spring with a stiffness *k*_*RH*_ = *πRP*, where *R* is the RH radius near the tip and *P* is the turgor pressure(31). The plot of *dL* as a function of 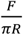 (figure 5B) shows a linear trend, consistent with the predicted behavior, with the slope corresponding to 1/*P*. From the linear fit, we estimated the mean turgor pressure value across the 41 RH tested to be *P* = 1,7.10^5^ *Pa* ± 0,5.10^5^ *Pa*. This turgor pressure estimate falls within the correct range of turgor pressure reported previously(32).

**Figure 5.**
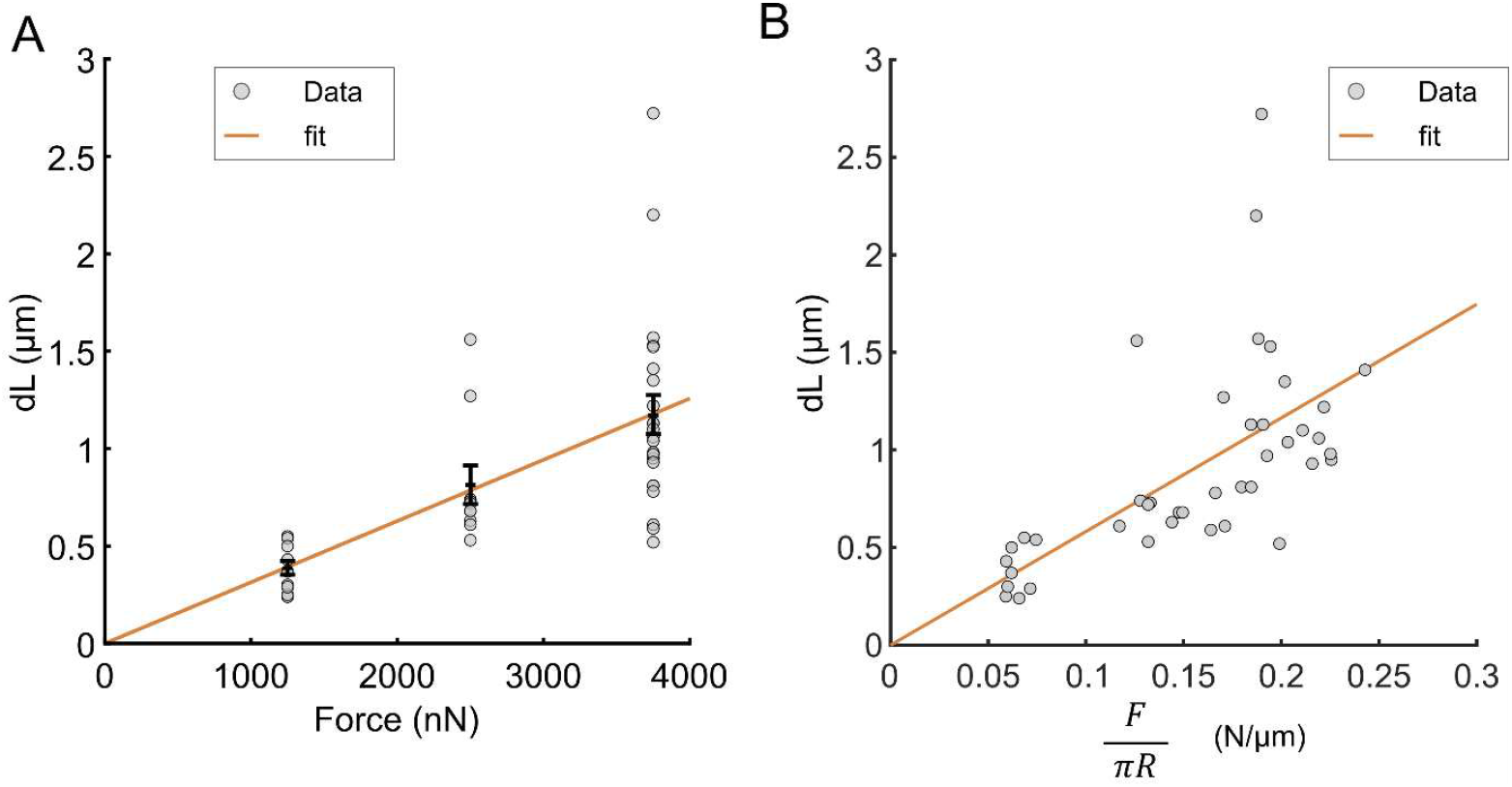
Instantaneous RH Compression upon axial force application. **A**. Root hair compression dL as a function of the applied force (n=41, N=12).The black error bars represent the average dL +/-sem for each force exerted. The orange line corresponds to a linear fit of all data points with the origin set at 0. **B**. Root hair compression dL as a function of F/πR (n=41). The orange line corresponds to a linear fit of all data points with the origin set at 0.

## Discussion

Adapting rheology techniques to quantify how an axial resisting force affects the growth of a single tip-growing root hair presents a significant challenge. These elongated, cylindrical outgrowths of epidermal root cells are highly fragile and prone to buckling under axial compression (27). To address this, our experimental setup was specifically designed to prevent physical buckling while applying a controlled force sufficient to reduce the growth rate.

To ensure stable anchoring of the root hair base during force application, roots were cultivated in agar gels. For the experiments, agar was selectively removed from one side of the root, allowing force to be applied to the tip of newly emerging root hairs while keeping the root securely in place. The obstacle was precisely positioned orthogonally to the root hair, ensuring that the compressive force applied to the tip was indeed axial.

The force was maintained at a constant level below the buckling threshold using a feedback loop that adjusted the position of the entire root, keeping the root hair tip stationary as it grew. The main limitations of our approach are the spatial (<60 nm) and temporal (0.1 s) resolutions of the obstacle’s position, which result in a precision of less than 1% for the constant force value.

Notably, since the force is kept constant during root hair growth, there is no elastic contribution to changes in root hair length, except for the very brief period immediately following the application of force (<4 s). This observation justifies the use of a viscoplastic model to describe root hair growth, specifically in terms of cell wall flow and elongation.

The growth rate was observed to decrease as a function of both the applied force (Fig. 3C) and the resulting stress generated in the root hair wall, 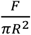 (Fig 3.D). When the data were fitted with a linearly decreasing function, the fit quality improved significantly when using 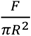 as the variable (p-value = 2.16 × 10^−19^ for Fig. 3D, compared to 1.7 × 10^−16^ for Fig. 3C). This result is consistent with Equation (3), which corresponds to Lockhart’s model(15) as adapted by Minc *et al*.(3) to describe the feedback of force on apical growth in fission yeast. Equation (3) relates growth to cell wall deformation, assuming that growth is not limited by water entry, a condition consistent with the characteristics of the root hairs examined in this study (Section 2 of the Supplementary Material). Under these conditions, the linear fit of the velocity data as a function of the stress applied by the obstacle 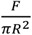 (Figure 3D) makes it possible, using the values of the slope and the x-intercept at zero velocity, to provide an initial estimate of the extensibility (*φ*) and the excess pressure (*P* − *Y*_*p*_), respectively.

However, these initial estimates are interdependent due to the fitting process and are sensitive to the variability in RH behaviour, as evidenced by the dispersion in growth rate values at zero applied force (Fig. 3D). To obtain independent estimates of the excess pressure (*P* − *Y*_*p*_) and extensibility *φ*, we relied on Equations (4) and (5), which express the ratio 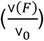 and difference (V_0_ – V(*F*)) between the measured rates of RH growth before and after force application. As shown in Figure 4, this approach significantly reduced the variability in the data and, consequently, minimized the uncertainty in the estimated parameters.

The low dispersion of the data for the ratio and difference of growth rates before and after force application around the lines corresponding to the linear fit (Figures 4A and B) is remarkable in two respects. First, it confirms the validity of Lockhart’s model for tip growth, which has been rigorously validated only a few times(20) and from which the excess pressure above the yield threshold and the extensibility parameters have never been quantified before(5). Second, the fact that the large number of data in Figure 4 (51 different root hairs from 13 different plants) follow so well the linear fit suggests that the parameters are conserved form cell to cell and that they are strongly regulated by the plant. This is particularly true for the excess pressure, given how limited the dispersion around the linear fit is (Figure 4A).

Finally, the measured instantaneous compression of the RH tip upon force application enabled us to estimate RH turgor pressure (1,7.10^5^ *Pa*) leading to an estimated yield turgor pressure of *Y*_*p*_∼0,7.10^5^ *Pa*. This value aligns closely with our previous estimation(27). This result is particularly robust, as both the excess pressure and turgor pressure were independently measured on each cell during a single experiment.

Note that the value of the turgor pressure (*P*∼1,7.10^5^ *Pa*), and consequently the threshold value (*Y*_*p*_∼0,7.10^5^ *Pa*.), are certainly underestimated. Indeed, to estimate *P*, we used a Hertz-type model, assuming that the tip of the root hair was a spherical cap with a radius equal to that of the root hair itself. However, the tip is certainly not spherical, and the local contact radius with the obstacle is likely smaller than the radius of the root hair. Since the expression for the RH stiffness *k*_*RH*_ = *πRP* implies a higher pressure for a smaller radius, this leads to an underestimation.

As a perspective, our setup provides a robust and reproducible quantitative mean to test the invasive ability of tip growing cells and their response to environmental cues. The technique presented in this article can be adapted to most tip-growing species. It could provide quantitative insight on turgor and cell wall modulation that can complement and explain previously observed morphological changes in response to environmental cues(1, 8, 33). Regarding root hairs, the ability to cope with a force while growing and, more importantly, turgor and cell wall adaptation can now be tested under various soil conditions that are known to affect root hair growth such as low phosphate(34, 35) and hypertonic mediums(36). The role of cellular parameters on this invasive ability can be further deciphered by phenotyping mutants (cell wall elasticity, turgor regulation). Moreover, the compatibility of our setup with fluorescence microscopy enables tracking intracellular biofluorescent markers upon force application. This could provide a groundbreaking tool for investigating mechanosensitive pathways involved in root hair tip growth.

## Materials and Methods

### Device preparation

The microfluidic-like system(29) used to grow the roots was made with a custom-made mould of PVC with a single channel of 270 µm height and 1cm wide. The PDMS 1/10 base/curing agent mixture (Sylgard 184, Dow Corning) is spilled on the mould. The ensemble was then cured at 65°C overnight. The bounding of the PDMS chip with the glass microscope slide was then done with a plasma cleaner (Harrick Plasma, PDC-002-CE). The channel was then filled with ½ MS (MS Murashige and Skoog) medium with 5% sucrose (w/w) and 1% agar (w/w) (Duchefa, plant agar). Before filling the channel, the PH of the medium was adjusted to 5.7. A 0.5cm thick layer of the same medium was poured all around the PDMS chip.

### Plant culture

Arabidopsis seeds expressing a fluorescent inner nuclear membrane protein pSUN1:SUN1-GFP(37) were used. The seeds were sterilized and then stratified in an Eppendorf filled with 1mL of ½ MS (MS Murashige and Skoog) medium with 5% sucrose (w/w) for 48h at 5°C. After stratification, the seeds were deposited on the agar-filled device close to the entrance of the channel. The microfluidic-like system was then placed in a petri dish sealed with microporous tape. The dish was then positioned at a 45° angle with respect to the vertical in an incubator (Sanyo, versatile environmental test chamber MLR-351H) with an incubation cycle of 16h light, 20.5°c and 8h dark, 17°C, at 65% humidity.

### Cantilever preparation and calibration

The microplates were made by stretching (Narishige PB-7 puller, Japan) a glass plate cutting it and fusing it with a glass capillary as described before(25). The calibration of the glass microplate was done as described before(25) by calibrating it with a standard glass microplate.

### Force application and data collection

Before the experiment the plant and the microfluidic-like system were placed on a microscope holder. Using tweezers and a scalpel, the agar on one side of the root tip was removed and replaced with ½ MS (MS Murashige and Skoog) liquid medium with 5% sucrose (w/w), PH 5.7. The system was then placed under an IX83 Olympus microscope and observed with a 40 × 0.55 NA objective. The microplate was then placed in position using a 3D micromanipulator system to ensure that its position can be recorded by a position sensitive detector (S3979 one-dimensional Position Sensitive Detector Hamamatsu, France). The root was then placed at an angle such that one root hair grew against the microplate. The feedback loop was adapted from a previously described experiment(28). The position of the piezoelectric stage was recorded during the experiment and bright field images were taken to assess the root hair growth direction. The measured standard deviation of the microplate position (measured by the PSD detector) was smaller than 60 nm, which corresponds to a precision on the applied force better than 1% of the target force. In all experiments, growth speed was estimated by fitting the RH tip position versus time as measured from the feedback loop control signal acting on the piezoelectric stage displacement, with a confidence interval of ∼2.10^-4^ µm/min.

### Statistical analysis

The Data were analysed using Matlab. The data are presented as mean values +/-s.e.m. For the values determined by a fit the Matlab Curve Fitting tool was used and data are presented as fitted value +/-half-width of the 95% confidence interval.

## Acknowledgments

The study was partially supported by the Université Paris Cité, Idex ANR-18-IDEX-0001, funded by the French Government through its “Investments for the Future” program, and also by the projects “Mecha-Nuc” ANR-20CE13-0025-03 and “scEm-bryoMech” ANR-21-CE13-0046. PDS acknowledges support from HFSPO grant number 2022-RG107.

## Notes

### Competing Interest Statement

The authors have declared no competing interest.

## References

1. A. Sanati Nezhad, M. Naghavi, M. Packirisamy, R. Bhat, A. Geitmann, Quantification of cellular penetrative forces using lab-on-a-chip technology and finite element modeling. Proc. Natl. Acad. Sci. 110, 8093–8098 (2013).

2. C. M. Rounds, M. Bezanilla, Growth Mechanisms in Tip-Growing Plant Cells. Annu. Rev. Plant Biol. 64, 243–265 (2013).

3. N. Minc, A. Boudaoud, F. Chang, Mechanical Forces of Fission Yeast Growth. Curr. Biol. 19, 1096–1101 (2009).

4. N. P. Money, Insights on the mechanics of hyphal growth. Fungal Biol. Rev. 22, 71–76 (2008).

5. J. Dumais, Mechanics and hydraulics of pollen tube growth. New Phytol. 232, 1549–1565 (2021).

6. S. L. Shaw, J. Dumais, S. R. Long, Cell Surface Expansion in Polarly Growing Root Hairs of Medicago truncatula. Plant Physiol. 124, 959–970 (2000).

7. J. T. Burri, et al., Feeling the force: how pollen tubes deal with obstacles. New Phytol. 220, 187–195 (2018).

8. D. D. Thomson, et al., Contact-induced apical asymmetry drives the thigmotropic responses of C andida albicans hyphae: Asymmetry drives thigmotropic responses. Cell. Microbiol. 17, 342–354 (2015).

9. N. Yanagisawa, N. Sugimoto, H. Arata, T. Higashiyama, Y. Sato, Capability of tip-growing plant cells to penetrate into extremely narrow gaps. Sci. Rep. 7, 1403 (2017).

10. M. Zarebanadkouki, P. Trtik, F. Hayat, A. Carminati, A. Kaestner, Root water uptake and its pathways across the root: quantification at the cellular scale. Sci. Rep. 9, 12979 (2019).

11. C. J. Stubbs, D. D. Cook, K. J. Niklas, A general review of the biomechanics of root anchorage. J. Exp. Bot. 70, 3439–3451 (2019).

12. C. Puerner, et al., Mechanical force-induced morphology changes in a human fungal pathogen. BMC Biol. 18, 122 (2020).

13. R. Reimann, et al., Durotropic Growth of Pollen Tubes. Plant Physiol. 183, 558–569 (2020).

14. D. Pereira, T. Alline, L. Cascaro, E. Lin, A. Asnacios, Mechanical resistance of the environment affects root hair growth and nucleus dynamics. Sci. Rep. 14, 13788 (2024).

15. J. A. Lockhart, An analysis of irreversible plant cell elongation. J. Theor. Biol. 8, 264–275 (1965).

16. M. Quiros, M.-B. Bogeat-Triboulot, E. Couturier, E. Kolb, Plant root growth against a mechanical obstacle: the early growth response of a maize root facing an axial resistance is consistent with the Lockhart model. J. R. Soc. Interface 19, 20220266 (2022).

17. A. E. Hill, B. Shachar-Hill, J. N. Skepper, J. Powell, Y. Shachar-Hill, An Osmotic Model of the Growing Pollen Tube. PLoS ONE 7, e36585 (2012).

18. J. F. Abenza, et al., Wall mechanics and exocytosis define the shape of growth domains in fission yeast. Nat. Commun. 6, 8400 (2015).

19. E. R. Rojas, S. Hotton, J. Dumais, Chemically Mediated Mechanical Expansion of the Pollen Tube Cell Wall. Biophys. J. 101, 1844–1853 (2011).

20. R. Benkert, G. Obermeyer, F.-W. Bentrup, The turgor pressure of growing lily pollen tubes. Protoplasma 198, 1–8 (1997).

21. D. Hüsken, E. Steudle, U. Zimmermann, Pressure Probe Technique for Measuring Water Relations of Cells in Higher Plants. Plant Physiol. 61, 158–163 (1978).

22. D. Suslov, K. Vissenberg, “Cell Wall Expansion as Viewed by the Creep Method” in Plant Biomechanics, A. Geitmann, J. Gril, Eds. (Springer International Publishing, 2018), pp. 305–320.

23. D. J. Cosgrove, Wall relaxation in growing stems: comparison of four species and assessment of measurement techniques. Planta 171, 266–278 (1987).

24. N. Desprat, A. Richert, J. Simeon, A. Asnacios, Creep Function of a Single Living Cell. Biophys. J. 88, 2224–2233 (2005).

25. N. Desprat, A. Guiroy, A. Asnacios, Microplates-based rheometer for a single living cell. Rev. Sci. Instrum. 77, 055111 (2006).

26. S. Robinson, C. Kuhlemeier, Global Compression Reorients Cortical Microtubules in Arabidopsis Hypocotyl Epidermis and Promotes Growth. Curr. Biol. 28, 1794–1802.e2 (2018).

27. T. Alline, L. Cascaro, D. Pereira, A. Asnacios, Micro-mechanical approaches to characterize tip growth: Insights into Root Hair Elasto-Viscoplastic Properties. [Preprint] (2025). Available at: https://www.biorxiv.org/content/10.1101/2025.08.06.668718v1 [Accessed 8 August 2025].

28. D. Mitrossilis, et al., Real-time single-cell response to stiffness. Proc. Natl. Acad. Sci. 107, 16518–16523 (2010).

29. D. Pereira, T. Alline, G. Singh, M.-E. Chabouté, A. Asnacios, “A Microfluidic-Like System (MLS) to Grow, Image, and Quantitatively Characterize Rigidity Sensing by Plant’s Roots and Root Hair Cells” in Mechanobiology, Methods in Molecular Biology., R. Zaidel-Bar, Ed. (Springer US, 2023), pp. 121–131.

30. M. Akkerman, et al., Texture of cellulose microfibrils of root hair cell walls of Arabidopsis thaliana, Medicago truncatula, and Vicia sativa. J. Microsc. 247, 60–67 (2012).

31. P. Durand-Smet, E. Gauquelin, N. Chastrette, A. Boudaoud, A. Asnacios, Estimation of turgor pressure through comparison between single plant cell and pressurized shell mechanics. Phys. Biol. 14, 055002 (2017).

32. C. Municio-Diaz, et al., Mechanobiology of the cell wall – insights from tip-growing plant and fungal cells. J. Cell Sci. 135, jcs259208 (2022).

33. X. Zhou, et al., Membrane receptor-mediated mechano-transduction maintains cell integrity during pollen tube growth within the pistil. Dev. Cell 56, 1030–1042.e6 (2021).

34. S. Datta, H. Prescott, L. Dolan, Intensity of a pulse of RSL4 transcription factor synthesis determines Arabidopsis root hair cell size. Nat. Plants 1, 15138 (2015).

35. T. R. Bates, J. P. Lynch, Stimulation of root hair elongation in Arabidopsis thaliana by low phosphorus availability. Plant Cell Environ. 19, 529–538 (1996).

36. M. Volgger, I. Lang, M. Ovečka, I. Lichtscheidl, Plasmolysis and cell wall deposition in wheat root hairs under osmotic stress. Protoplasma 243, 51–62 (2010).

37. K. Graumann, J. Runions, D. E. Evans, Characterization of SUN-domain proteins at the higher plant nuclear envelope. Plant J. 61, 134–144 (2010).

