## Supplemental Material for "Single Root hair growth under constant force: insights into wall mechanics"

#### 1) Viscoplastic Model

The root hair is described as a cylinder of radius R and length L. The turgor pressure is considered constant and denoted P. We consider the cell wall as a homogeneous material with thickness h. At the tip, new cell wall material and cell wall remodelling factor fluxes allow for the root hair to grow longitudinally only at the tip in a region of length l.

The longitudinal stress in the wall due to the turgor pressure is noted

$$\sigma_L = \frac{PR}{2h} \qquad (S1)$$

The growth is described through a viscoplastic model. The longitudinal strain is proportional to the stress above yield threshold noted $Y_\sigma$.

$$\dot{\varepsilon}_L = \varphi(\sigma_L - Y_\sigma) \qquad (S2)$$

$\varphi$ is the extensibility, this quantity can be interpreted as an inverse viscosity. It describes the irreversible deformation of the cell wall under stress. It encompasses the probable cell wall remodelling. Here we neglected the geometry of the tip and the possible inhomogeneity of this quantity in the region of the tip. The extensibility is thus an effective extensibility. This formulation of the strain is a formulation of the Lockhart law in stress.

Since growth only occurs at the tip, we can relate the strain to the cell longitudinal growth $v_0$ as follow:

$$v_0 = \frac{dL}{dt} = l\,\varphi(\sigma_L - Y_\sigma) \qquad (S3)$$

This law can be written in pressure with a yield in pressure $Y_P = \frac{2hY_\sigma}{R}$ as follow

$$v_0 = \frac{lR}{2h}\,\varphi(P - Y_P) \qquad (S4)$$

When a longitudinal force opposing root hair growth is applied at the tip, the stress in the cell wall is reduced.

$$\sigma_L(F) = \frac{PR}{2h} - \frac{F}{2\pi Rh} \qquad (S5)$$

The new growth rate $v(F)$ following equation S2 is:

$$v(F) = l\varphi\left(\frac{PR}{2h} - \frac{F}{2\pi Rh} - Y_\sigma\right) \qquad (S6)$$

This translated in term of pressure gives:

$$v(F) = \frac{lR}{2h}\,\varphi\left(P - \frac{F}{\pi R^2} - Y_P\right) \qquad (S7)$$

This finally gives 2 relations between the growth speed before and after force application:

$$\frac{v(F)}{v_0} = \left(1 - \frac{F}{\pi R^2\left(P - Y_p\right)}\right) \qquad (S8)$$

$$v(F) - v_0 = \frac{l\varphi F}{2\pi hR} \qquad (S9)$$

### 2) Growth is limited by wall deformation

In this model we considered that the water fluxes do not limit growth. Here, we justify this hypothesis by roughly estimating how far from the osmotic equilibrium the RH are.

If we consider the Lockhart equation describing the water entry in the cell[1] and thus if we consider the cell as a perfect osmometer[2], we get:

$$\mathrm{v}(t) = L_p \frac{2L}{R}(\Delta\Pi - P) \qquad (S.10)$$

$\Delta\Pi$ is the osmotic potential difference between the interior and the exterior of the cell. $L_p$ is the plasma membrane permeability. Classical values of $L_p$ range between $10^{-13}\ m.s^{-1}Pa^{-1}$ and $10^{-11}\ m.s^{-1}Pa^{-1}$ depending on the presence of aquaporin channels[3]. We can thus take a low value estimate of $L_p = 10^{-13}\ m.s^{-1}Pa^{-1}$ to estimate a maximal value for $(\Delta\Pi - P)$:

$$(\Delta\Pi - P) = \frac{\mathrm{v}R}{2L_pL} \qquad (S.11)$$

By taking $R = 5\mu m$, $L = 100\ \mu m$ and $\mathrm{v} = 1{,}2\ \mu m.min^{-1}$, which is the measured value for root hair free growth speed in fig 3.C of the main text, we get $(\Delta\Pi - P) = 5000\ Pa \ll P \sim 2.10^5\ Pa$. Thus $P \approx \Delta\Pi$, and the cell is always very close to the osmotic equilibrium. This corresponds to the growth regime in which cell wall deformation limits growth[4].

### References


1. Lockhart, J. A. An analysis of irreversible plant cell elongation. *Journal of Theoretical Biology* **8**, 264–275 (1965).
2. Dainty, J. Water Relations of Plant Cells.
3. Forterre, Y. Chapter 1. Basic Soft Matter for Plants. in *Soft Matter Series* (eds. Jensen, K. & Forterre, Y.) 1–65 (Royal Society of Chemistry, Cambridge, 2022). doi:10.1039/9781839161162-00001.
4. Dumais, J. Mechanics and hydraulics of pollen tube growth. *New Phytol* **232**, 1549–1565 (2021).
